# Pneumatic Controlled Nano-Sieve for Efficient Capture and Release of Nanoparticles

**DOI:** 10.1101/2022.09.30.510404

**Authors:** Animesh Nanaware, Taylor Kranbuhl, Jesus Ching, Janice S. Chen, Xinye Chen, Qingsong Tu, Ke Du

**Affiliations:** Department of Mechanical Engineering, Rochester Institute of Technology, Rochester, NY 14623, USA; Department of Chemical Engineering, Rochester Institute of Technology, Rochester, NY 14623, USA; Mammoth Biosciences, Inc., Brisbane, California, USA; Department of Chemical and Environmental Engineering, University of California, Riverside, CA 92521, USA

## Abstract

A pneumatic controlled nano-sieve device is demonstrated for the efficient capture and release of 15 nm quantum dots. This device consists of a 200 nm deep glass channel and a PDMS-based pneumatic pressure layer to enhance target capture. The fluid motion inside the nano-sieve is studied by computational fluidic dynamics (CFD) and microfluidic experiments, enabling efficient target capture with a flow rate as high as 100 μL/min. In addition, micro-grooves are fabricated inside the nano-sieve to create low flow rate regions, which further improves the target capture efficiency. A velocity contour plot is constructed with CFD, revealing the flow rate is lowest at the top and bottom of the micro-grooves. This phenomenon is supported by the observed nanoparticle clusters surrounding the micro-grooves. By changing the morphology and pneumatic pressure, this device will also facilitate rapid capture and release of various biomolecules.

## I. INTRODUCTION

The efficient isolation and retrieval of nanostructures and biomolecules are the key for numerous biomedical applications such as molecular diagnostics^1,2^, precision therapeutics^3,4^, and mechanobiology^5,6^. Nanofiltration membranes with controllable selectivity emerged boasting the advantages of being low cost and easy to use^7,8^. However, post-treatments are always needed to remove the captured targets which are complex and can alter the properties of the membrane^9^. On the other hand, electrophoresis has been developed to separate nanoparticles^10,11^, nucleic acids^12,13^, proteins^14,15^, and cells^16,17^. However, a high DC voltage is required for the separation, and it cannot be directly used in the complicated biofluids as it can change the electrostatic force in the solution.

Various micro- and nanofluidic systems have been created in recent years with the advantages of high scalability and resolution^16,17^. A simple confinement approach by a convex lens and a flat substrate has been introduced to better trap molecules for fluorescence sensing^18–20^. However, only one single trapped region can be formed by this approach. On the other hand, top-down approaches can pattern delicate nanofeatures with controllable geometries in the nanofluidic channel, such as dual height and staircase nanochannels^21–23^. However, the fabrication and bonding of these rigid chips are challenging. Recently, Wunsch et. al. created a nanoscale lateral displacement chip with periodic nanopillars to separate exosomes from colloids with a size down to 20 nm^24^. It has also been used to manipulate and stretch lamda DNA in the 2-D nanopatterns^25^. Further, Yu et. al. demonstrated single-cell capture and single-DNA linearization with a pneumatic microchannel and nanoslits, thus paving the ways for a fully on-chip, nucleic acid extraction and genome sequencing device^26^.

Recently, we developed a nano-sieve device for the efficient capture of microparticles by controlling the hydrodynamic pressure in a deformable channel^27^. By trapping different sizes of microparticles, this device has been used for the separation, concentration, and retrieval of microorganisms and red blood cells from bodily fluids at a high flow rate, designed for on-chip sepsis diagnosis^28–30^. Even though microparticles can be efficiently captured and released, this device cannot be used to capture nanoparticles (smaller than 100 nm) as the channel height will be too high to capture the small nanoparticles.

In this work, we present an improved nano-sieve device for the capture and release of quantum dots with a size of 15 nm. A pneumatic chamber is added on top of the nano-sieve to counterbalance the hydrodynamic pressure produced in the channel by the high flow rate. With a pneumatic pressure of 14 psi, a high capture efficiency is achieved with a high flow rate of 100 μL/min. A numerical model based on computational fluid dynamics (CFD) is developed to analyze the characteristics of the fluid motion and the distribution of the hydrodynamic pressure within the channel. In addition, guided by CFD, micro-grooves with a feature size of ~5 μm are designed and patterned in the nano-sieve to further increase the capture efficiency. Combined with our numerical simulation and experimental results, we show that the micro-grooves reduce the flow rate around them thus enhancing the trapping of the quantum dots. Lastly, the captured targets can easily be retrieved for downstream analysis by turning off the positive pneumatic pressure.

## II. EXPERIMENTAL

### A. Nano-sieve fabrication

A 200 nm Tetraethyl Orthosilicate (TEOS) layer was deposited on a 6-inch glass wafer with plasma-enhanced chemical vapor deposition PECVD (AMAT P5000). Then, a ~1.05 μm thick positive photoresist (HMDS prime) was deposited on the TEOS layer with a SVG track. An ASML Stepper (field size – 22 × 22mm, NA – 0.48-0.6, 0.35 μm size capability) was used to define the channel and the microstructures. The dimension of each channel is 4 mm in width and 6 mm in length. After exposure, a CD-26 developer was used to remove the positive photoresist. To define the nano-sieve with TEOS, wet etching was performed with Buffer Oxide Etch - 10:1 (BOE) for 104 s. After wet etch, a Trion Asher was used to clean the channel. Finally, a sacrificial positive photoresist layer with a size of 4 mm × 6 mm was patterned on top of the nano-sieve channel with the ASML Stepper system.

### B. Pneumatic layer fabrication

To fabricate the air chamber, Polydimethylsiloxane (PDMS) base and curing agent (SYLGARD 184 silicone elastomer) were mixed in a weighted ratio of 10:1, followed by desiccating and pouring into a 3-D printed mold (See supplement material at URL). For forming the pneumatic membrane, another 10 mL of mixed PDMS was spin coated on a salinized glass wafer at 350 rpm for 1 min, giving an even thickness of 350 μm. The mold and the wafer were both baked in an oven at 40 °C for 4 hours. After baking, the pneumatic inlet was created with a 1 mm biopsy punch on the air chamber.

Plasma treatment was performed on the air chamber and the pneumatic membrane for 45 s each and pressed together, followed by baking it in an oven at 140 °C for 1 hour. After the bake step, the bonded pneumatic layer was peeled off from the 3-D printed mold, followed by creating a liquid inlet/outlet with a 1 mm biopsy punch. Final bonding was carried out between the nano-sieve and the pneumatic layer with plasma treatment for 45 s on each surface and with a final bake at 140 °C for 1 hour in an oven. Finally, the sacrificial photoresist layer was dissolved by filling acetone into the liquid channel.

### C. Fluorescence quantification

Quantum dots (Qdot^™^ 605 Streptavidin Conjugate) with a size of 15 nm are used as a target. A customized fluorometer with a 488 nm laser, a parabolic mirror, and a USB spectrometer (Ocean Optics 2000+) were used to quantify the fluorescence intensity of the quantum dots^31,32^. The laser power was set at 3 mW with an integration time of 3 s and scans to an average of 2. Each liquid sample (10 μL) was added into a microplate and aligned with the laser beam for the measurement.

### D. SEM imaging

Tescan Mira3 was used to characterize the nano-sieve device. To reduce the SEM charging effect, a ~20 nm gold layer was coated onto the sample using a sputter tool (SPI-Module Sputter Coater). The SEM imaging was carried out at 20kV.

## III. MODELLING

Computational fluid dynamics (CFD) implemented in COMSOL were employed to systematically investigate the fluid motion and transport of quantum dots in the microfluidic channel^33^. Geometry of the models is based on the SEM images of the fabricated micro-channels. The fluid motion in the channel was analyzed under different model geometries, inlet flow rates, and applied pneumatic pressures perpendicular to the fluid direction.

## IV. RESULTS AND DISCUSSION

**Figure 1a** shows the working principle of the pneumatic controlled nano-sieved device. A PDMS based pneumatic chamber is assembled on top of the nano-sieve and controlled by a pneumatic pump (Precigenome - “PG-MFC-4CH”). The SEM image of the micro-grooves in the channel is shown in **Figure 1b** with a center-to-center distance of 6 μm. The quantum dot solution was pumped into the device by a syringe pump (NE-300 “Just Infusion”) with various flow rates. The micro-grooves are used to create dead volume around them for nanoparticle trapping (**Figure 1c**). In the presence of positive pressure, the gap between the pneumatic membrane and the substrate decreases thus can better capture the targets, such as the applied quantum dots as shown in **Figure 1d**. The nano-sieve after target capture shown in **Figure 1e-i** shows the strong pink fluorescence, indicating the presence of quantum dots. After target release, the channel becomes clean again as the quantum dots are washed to the outlet (**Figure 1e-ii**). The fabrication process of the pneumatic controlled nano-sieve is shown in **Figure 1f**: A thin TEOS layer (blue) is deposited on the glass substrate, followed by the patterning of photoresist (yellow). The TEOS layer is wet etched, followed by the patterning of the sacrificial photoresist. The pneumatic layer is created by bonding a pneumatic hollow chamber and a pneumatic membrane (purple). After bonding the chamber and the membrane, the entire pneumatic layer is then transferred and bonded onto the glass-based nano-sieve device.

**Figure 1.**
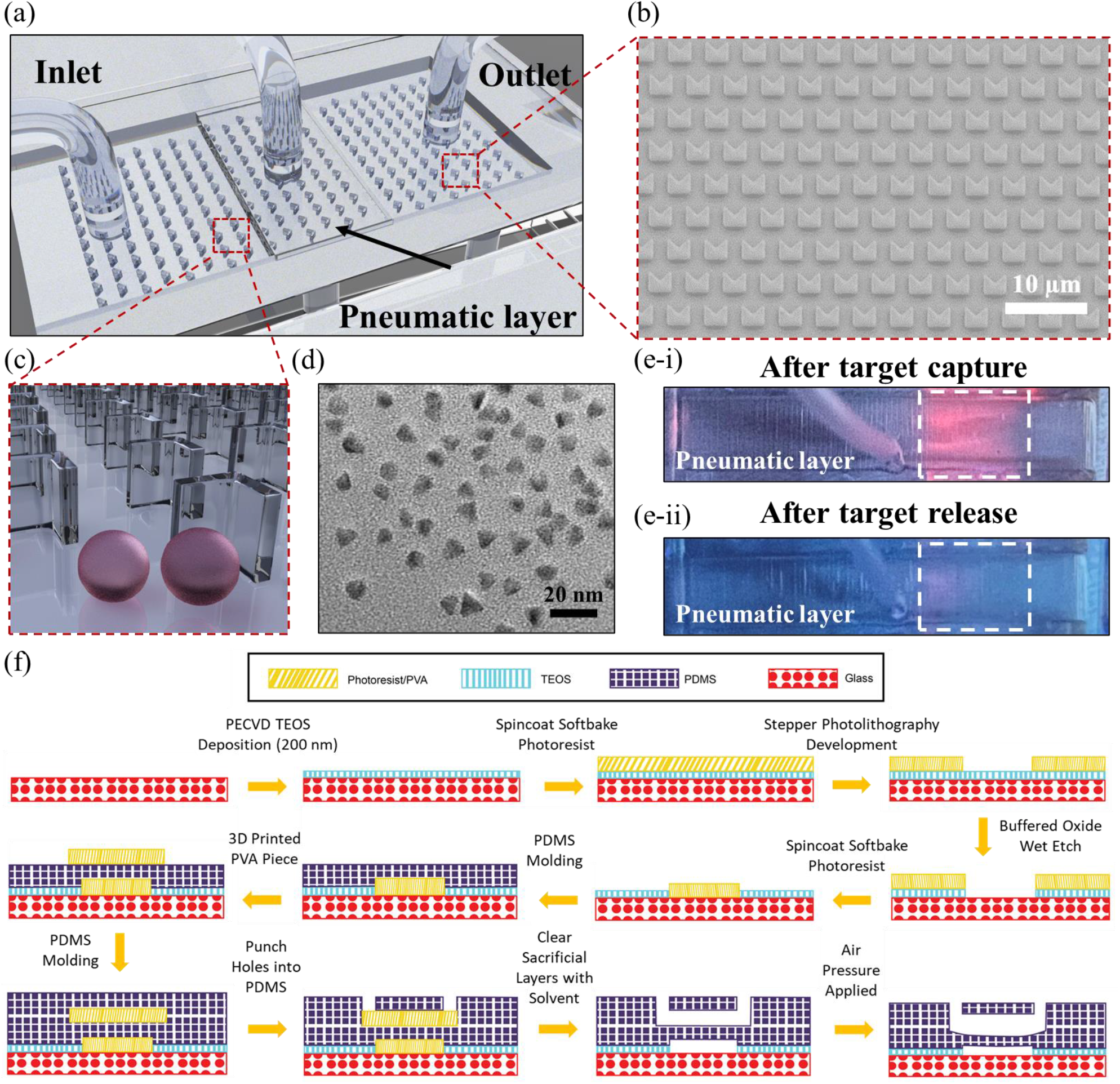
(a) Schematic of the pneumatic controlled nano-sieve device. (b) SEM image of the micro-grooves inside the channel with a feature size of 5 μm. (c) Schematic of the micro-grooves patterned on the glass substrate with the introduced nanoparticles. (d) TEM image of the applied quantum dots. (e) Photograph of the channel: after target capture (e-i) and after target release (e-ii). (f) Fabrication flow of the device by bonding a pneumatic layer on glass substrate.

To study the effect of pneumatic pressure on target capture, we applied a pneumatic pressure ranging from 0 to 14 psi on a planar nano-sieve device (without micro-grooves) while changing the flow rate from 25 to 100 μL/min. For each measurement, 10 mL of the quantum dot solution (1 nM) was pumped into the device through the inlet and the supernatant was collected at the outlet. A lower fluorescence intensity indicates a higher capture efficiency as less quantum dots are leaked through. As shown in **Figure 2a**, without positive pneumatic pressure, more quantum dots passed through the device uncaptured. This was observed by a fluorescent peak intensity reaching ~6,500 counts (green). The leaking is decreased with lower flow rate, as the hydrodynamic deformation was smaller at lower flow rates. Increasing the pneumatic pressure to 5 psi reduces target leaking. As shown in **Figure 2b**, the peak intensity for 100 μL/min sample drops to ~5,000 counts, which is lower than the 0 psi case. We further increased the pneumatic pressure to 10 psi and found that the peak intensity of the 100 μL/min sample drops to ~3,500 counts (**Figure 2c**). The 75 μL/min sample (blue) shows a weak signal, at lower flow rates, as no distinct peak was observed. With 14 psi pneumatic pressure (**Figure 2d**), only a weak peak is observed for both the 75 μL/min and 100 μL/min samples, indicating most of the quantum dots were capture within the device. The highest pneumatic pressure used in this study is 14 psi as higher pressure causes backflow of the sample solution.

**Figure 2.**
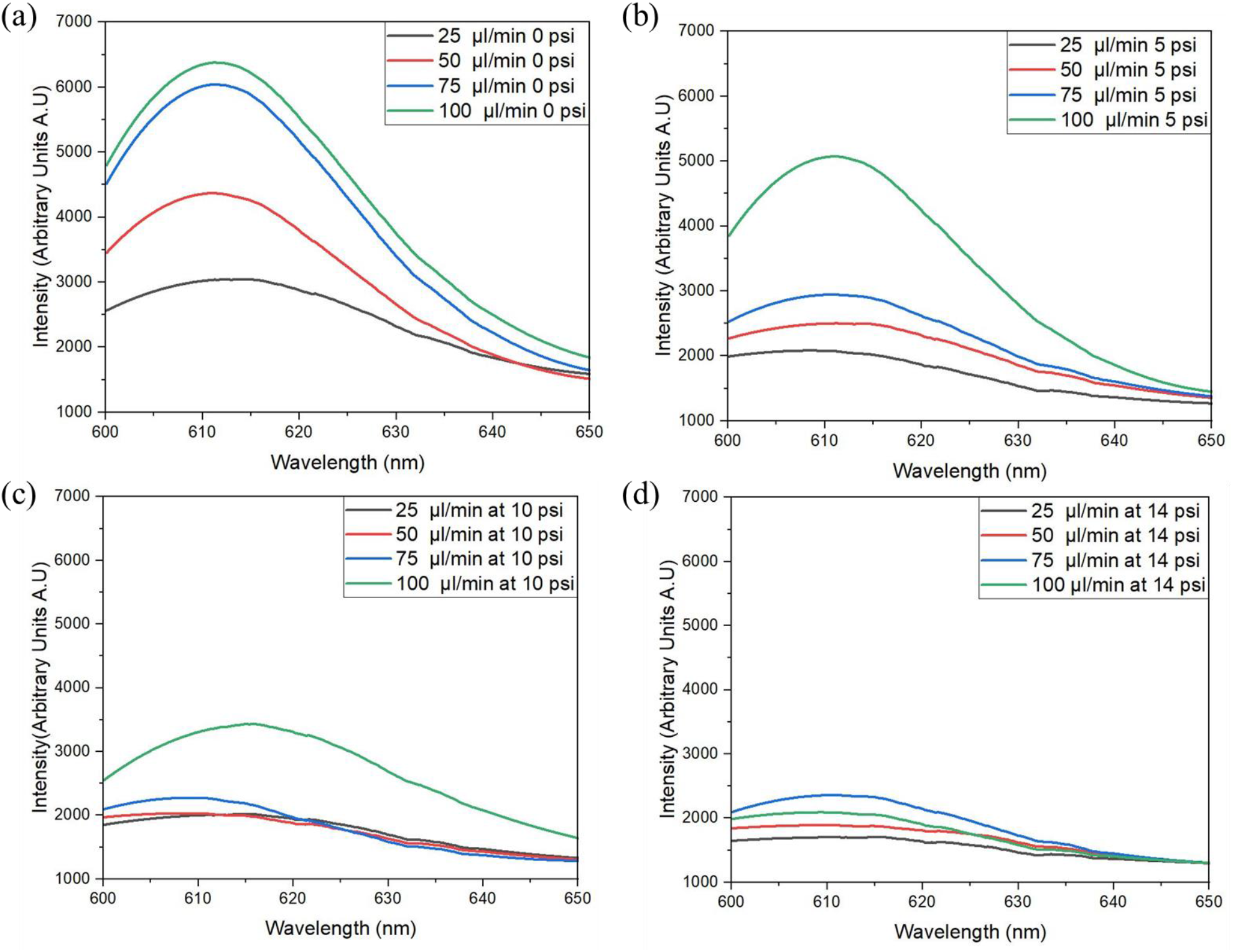
Uncorrected emission signal of the collected quantum dot solution from the nano-sieve outlet with a pneumatic pressure of (a) 0 psi, (b) 5 psi, (c) 10 psi, (d) 14 psi. Flow rate ranging from 25 μL/min to 100 μL/min is tested at each pneumatic pressure.

It is clear that the performance of quantum dot capture was affected by the inlet flow rate and the applied pneumatic pressure. To quantify their relations and optimize these parameters, we systematically studied their effects by modeling the fluid motion inside the microchannel with computational fluid dynamics (CFD). **Figure 3a** shows the schematics for the fluid motion in a single micro-groove microfluidics device cross-section. Constant pneumatic pressure was applied to the top of the microfluidic device, pushing down and deforming the membrane in the open channel. The fluid flow creates a hydrodynamic pressure that pushes back upward against the membrane. The greater the fluid flow is compressed within the channel, the higher the resultant hydrodynamic pressure force will be enacting on the membrane. The developed pressure near the microchannel under the flow rate of 0.15 mL/min is shown with the colored contour plots in **Figure 3b**. **Figure 3c** shows the pressure value near the entrance of the channel (black dashed line in Figure 3b) at three different flow rates. The pressure perturbation in the channel along the radial direction increases at higher flow rate due to the higher instability developed when the fluid motion is high; however, this perturbation is still very small (<5%) even at the highest flow rate of 1mL/min**. Figure 3d** shows the distribution of the developed pressure under several inlet flow rates, ranging from 0.1 μL/min to 1 mL/min.

**Figure 3.**
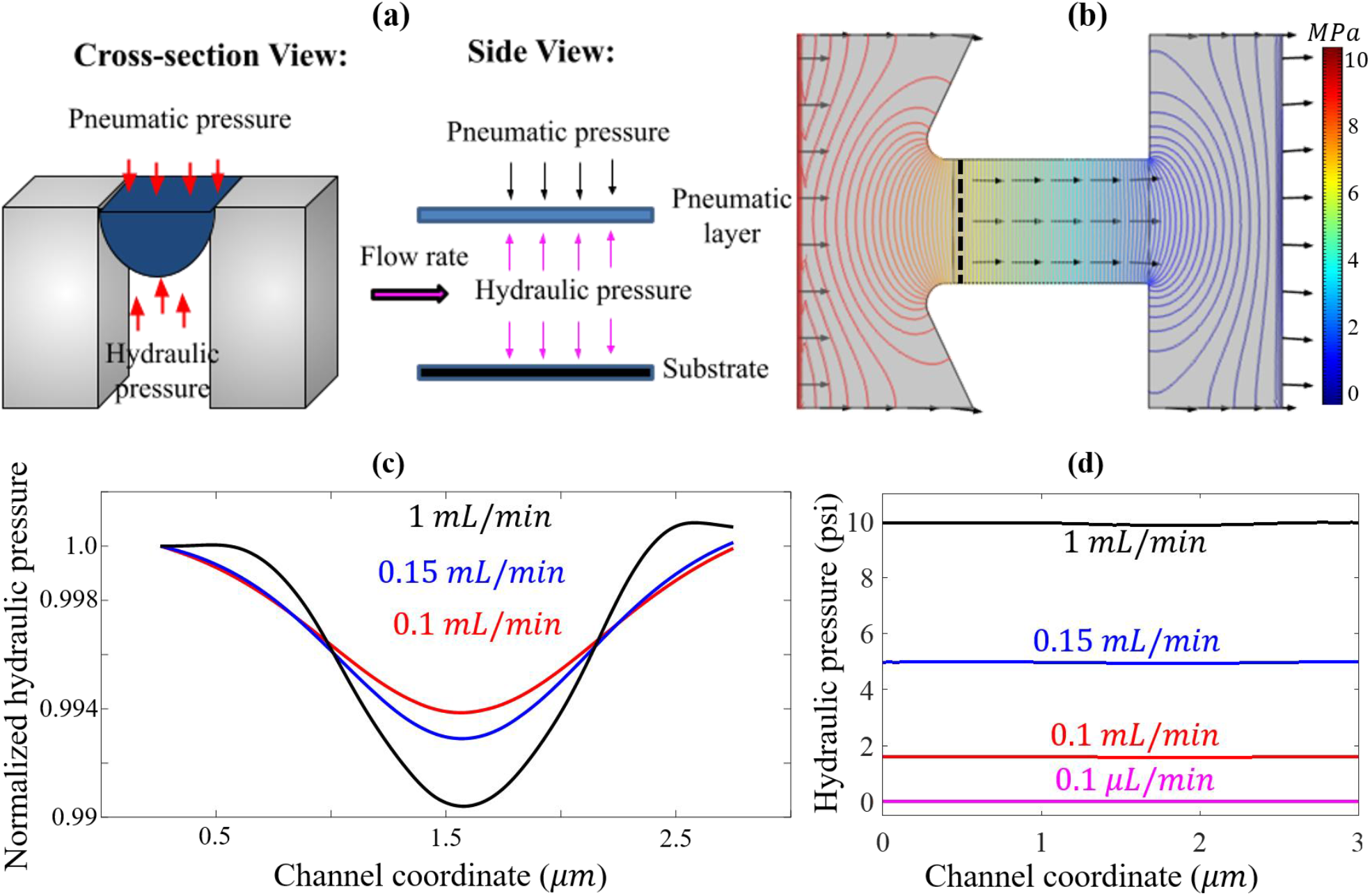
Modeling results to show the effects of pneumatic pressure on a microfluidics micro-groove. (a) Schematic cross-section and side view of a microfluidic micro-groove. b) Top view of hydraulic pressure of micro-groove when flow rate is 0.15 mL/min. c) Change in hydraulic pressure along the blue dash line in the channel in (b) under three different flow rates. The hydraulic pressure is normalized to the pressure value close to the wall of the channel. d) Hydraulic pressure along the blue dash line for several inlet flow rates.

We compared the target capture and release for flat and micro-grooves channels by using 5 psi pneumatic pressure and a 100 μL/min flow rate. For each measurement, the emission signal from 600 to 650 nm was integrated and the captured targets were counted by subtracting the collected supernatant and buffer background (see supplementary material at URL). As shown in **Figure 4a**, more quantum dots were trapped in the micro-groove channel with an average integrated signal of ~215,000 counts (green). On the other hand, the average integrated signal for the flat channel was only ~185,000 counts (orange). To retrieve the captured quantum dots, we injected 150 μL of buffer with a high flow rate (1,000 μL/min) to flush all the captured targets through the outlet. As shown in **Figure 4b**, there are more quantum dots being collected from the micro-groove channel (green) than the flat channel (orange), again proving that adding the microstructures can enhance the target capture. Interestingly, the capture efficiency for the flat channel had a greater variation than the micro-groove channel, which may be due to the instability in a wide and shallow channel at higher flow rates^34^.

**Figure 4.**
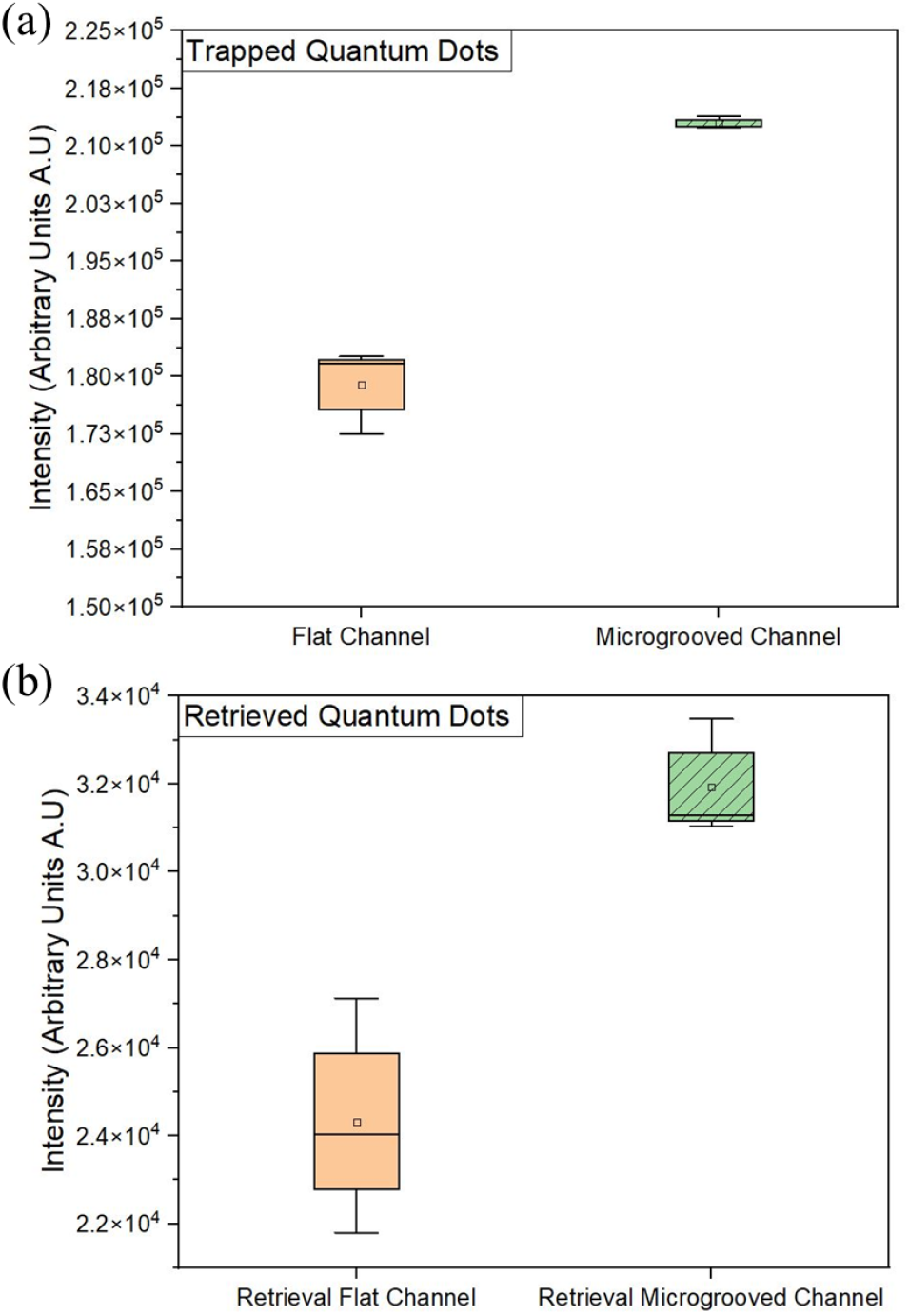
(a) Trapped and (b) Retrieved quantum dots for flat versus micro-groove channels. The pneumatic pressure and flow rate were set at 5 psi and 100 μL/min for all the samples. The fluorescence signal was integrated from 600 to 650 nm.

We combined SEM imaging and numerical simulation to explain why shallow micro-grooves can enhance target capture. To better visualize single particles, we used 100 nm microspheres (F8801 – Thermofisher Scientific) and injected the solution into the nano-sieve device at a flow rate of 25 μL/min. After that, the channel was washed using a buffer at a flow rate of 1000 μL/min. Finally, the pneumatic layer was carefully peeled-off and the channel was imaged by an SEM. As shown in **Figure 5a**, more nanoparticle clusters formed around the micro-grooves, especially surrounding at the bottom side. This was directly caused by the lower flow rate at these regions.

**Figure 5.**
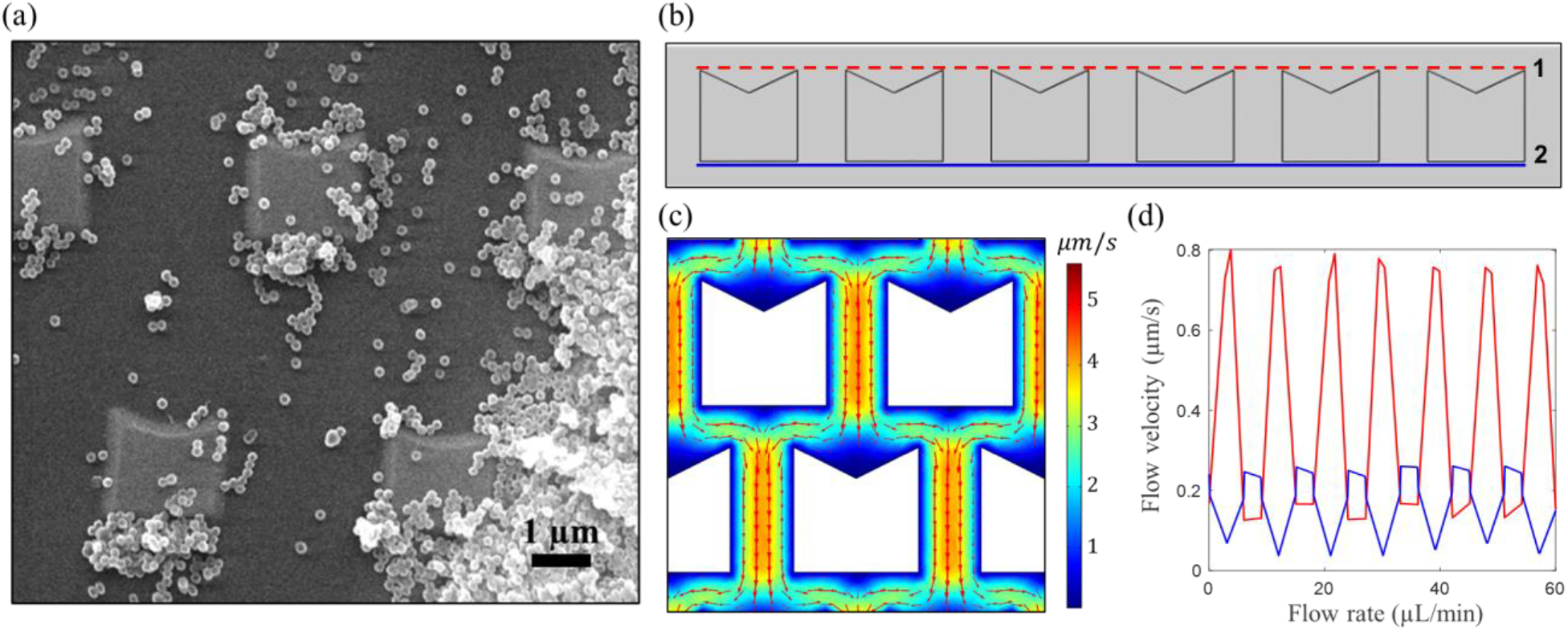
(a) SEM image to display nanosphere concentration around micro-grooves. (b) Top-view schematic of micro-grooves and cross sections at which velocity profiles were studied. (c) Velocity contour plot of several micro-grooves in the microfluidics device. (d) Velocity profile plots of the flow at the tip of the micro-groove and after the flow passes the micro-groove.

It is clear that more particles clustered at the top and bottom area of the micro-grooves. The rate of particle capture was directly related to the flow rate within the microfluidic device. The lower the flow rate, the higher the chance of particle capture will be since there is less turbulence forcing the particles to continue moving down the open channel. The velocity of the fluid moving in the channel was studied at the tip of the micro-groove and just as the flow passes the micro-groove, as shown in **Figure 5b**. In simulation, the flow velocity was lowest at the tips and bottom of the microstructure, which is shown in the contour plot **Figure 5c**. The darkest blue areas of the plot demonstrate that the flow velocity is almost 0 *μ*m/s. The velocity profile was taken along the x-coordinate arc length (**Figure 5d**), and the lowest velocity was aligned with the tips and bottom of the micro-groove.

Rapid isolation and concentration of small molecules from a large volume of solution is a key process for many clinical and environmental applications^35^. Instead of using surface chemistry method for separation, a pure isolation method is simpler and can completely avoid contamination issues. Electrokinetic flows have been developed for the efficient separation of different targets but cannot work with complicated bodily fluids as the electrostatic force will be altered. Therefore, front-end sample purification is always needed. Mechanical driven separation is simpler but it needs to counterbalance the hydrodynamic pressure built in the channel. The pneumatic controlled nano-sieve device developed by us has a very low aspect ratio (1:10,000) thus the hydrodynamic pressure is significantly reduced. A flexible pneumatic pressure layer is added to better capture and release the targets, which is more adaptable than rigid nanofluidic channels. Our device shows an excellent capture efficiency for small nanoparticles with a size of only ~15 nm. By applying a 14 psi positive pressure, the collected supernatant shows minimal fluorescence signal even with a flow rate as high as 100 μL/min. In addition, the captured nanoparticles can easily be retrieved by removing the positive pressure and flushing with a buffer solution. We further show that the micro-grooves can improve the capture efficiency by creating low flow rate regions near the structures. Similar strategies have been reported by other groups, but these all focus on larger targets such as single cells^36,37^. In the future, the morphology of the micro-grooves and surface property can be tuned to optimize the target capture and release.

One of the unique advantages of microfluidic-based isolation systems is the high multiplexing capability^38,39^. In our design, each channel is only 2 mm × 4 mm thus allowing the patterning of ~100 individual channels simultaneously. The flow rate and pneumatic pressure are regulated by digital pumps; thus, this device can work with microliter samples without causing errors. In the future, the multiplexing capability can be further improved by stacking the nano-sieve in a 3-D space^40^.

In this work, both quantum dots (15 nm) and microspheres (100 nm) were used. This shows that our system can work with various targets on the nanoscale, such as nucleic acids^41^, proteins^42^, antibodies^43^, and exosomes^44^. To separate multiple targets with different sizes in the same sample, a single channel can be designed with adaptable height, microstructure shape, and width. The mechanical properties of the deformable cap and the pneumatic pressure can also be tuned to work with even higher flow rates^45^. Established lysis and purification protocols can also be integrated in the nano-sieve to extract the desired targets^46^.

## ACKNOWLEDGMENTS

This research was supported by Mammoth Biosciences, Burroughs Wellcome Fund (1019955), and NSF CBET Project # 2035623. The authors would like to thank Mary Nguyen and Katherine Leising for the schematic design.

## DATA AVAILABILITY

Data available on request from the authors.

